# Chromatin loop anchors are associated with genome instability in cancer and recombination hotspots in the germline

**DOI:** 10.1101/166934

**Authors:** Vera B Kaiser, Colin A Semple

**Affiliations:** MRC Human Genetics Unit, MRC Institute of Genetics and Molecular Medicine, University of Edinburgh, Western General Hospital, Crewe Road, Edinburgh, EH4 2XU, UK

**Keywords:** cancer, tumour, recombination, structural variation, mutation, chromatin, Hi-C

## Abstract

Chromatin loops form a basic unit of interphase nuclear organisation, providing contacts between regulatory regions and target promoters, and forming higher level patterns defining self interacting domains. Recent studies have shown that mutations predicted to alter chromatin loops and domains are frequently observed in tumours and can result in the upregulation of oncogenes, but the combinations of selection and mutational bias underlying these observations remains unknown. Here, we explore the unusual mutational landscape associated with chromatin loop anchor points (LAPs), which are located at the base of chromatin loops and form a kinetic trap for cohesin. We show that LAPs are strongly depleted for single nucleotide variants (SNVs) in tumours, which is consistent with their relatively early replication timing. However, despite low SNV rates, LAPs emerge as sites of evolutionary innovation showing enrichment for structural variants (SVs). They harbour an excess of SV breakpoints in cancers, are prone to double strand breaks in somatic cells, and are bound by DNA repair complex proteins. Recurrently disrupted LAPs are often associated with genes annotated with functions in cell cycle transitions. An unexpectedly large fraction of LAPs (16%) also overlap known meiotic recombination hotspot (HSs), and are enriched for the core PRDM9 binding motif, suggesting that LAPs have been foci for diversity generated during recent human evolution. We suggest that the unusual chromatin structure at LAPs underlies the elevated SV rates observed, marking LAPs as sites of regulatory importance but also genomic fragility.

## INTRODUCTION

Recent evidence shows that many cancers and developmental disorders involve disruptions of chromatin organisation. Insertions and deletions are reported to alter the boundaries of topologically associating domains (TADs), which normally constrain the regulatory interactions of resident promoters and enhancers, causing dysregulated gene expression (Lupianez et al. 2016; Kaiser and Semple 2017). Disruptions of particular TAD boundaries have been reported in neuroblastoma (Peifer et al. 2015; Valentijn et al. 2015), medulloblastoma (Northcott et al. 2014), leukaemia (Groschel et al. 2014; Hnisz et al. 2016) and other cancers (Weischenfeldt et al. 2017), consistent with the hypothesis that structural variants (SVs) remodelling TAD boundaries may act as oncogenic ‘driver’ mutations under selection in tumour cells (Valton and Dekker 2016).

CTCF plays important roles in chromatin organisation, both demarcating domain boundaries as an insulator element (Dixon et al. 2012; Moore et al. 2015) and by bringing DNA sites that are distant in linear genomic distance intro close spatial proximity (Splinter et al. 2006). Pairs of CTCF binding sites may physically interact to form anchoring sites at the base of a chromatin loop, acting as physical barriers to the ring-shaped cohesion complex, which extrudes the loop (Haarhuis et al. 2017). Complex arrays of these loops are thought to make up the substructure of regulatory domains such as TADs (Rao et al. 2014), and recent experiments highlight the critical importance of CTCF for loop and TAD formation (Nora et al. 2017).

CTCF binding sites are highly mutated across cancer types, especially when they are located within loop anchor points (LAPs) (Katainen et al. 2015; Kaiser et al. 2016). Hyper-methylation of the GC rich CTCF binding motif has been shown to reduce CTCF binding in glioma, leading to the up-regulation of known oncogenes (Flavahan et al. 2016). Hnisz *et al.* (2016) have shown that constitutive CTCF-CTCF binding site interactions delineating loops are recurrently deleted in T-cell acute lymphoblastic leukemia, leading to oncogene activation. Overall, these data suggest that domain boundary or LAP lesions affecting gene regulation are far from rare in cancers, occurring at comparable rates to recurrent in-frame gene fusions (Weischenfeldt et al. 2017). However, it is unclear whether LAPs are intrinsically prone to high mutation rates in cancer, constituting a novel class of fragile sites in the genome, or whether the observed lesions affecting LAPs confer a selective advantage to tumour cells.

Somatic mutation rates vary across the genome, and a large fraction of this variation can be attributed to differences in replication timing, with late replicating regions of the genome accumulating increased levels of single nucleotide variants (SNVs) (Stamatoyannopoulos et al. 2009). Large regions of chromosomes (encompassing hundreds of Kb) are replicated synchronously in replication domains that correspond closely to TADs, linking chromatin organisation to spatiotemporal variation in replication (Dileep et al. 2015), while other, inter-correlated features of chromatin, such as histone methylation or acetylation patterns, are also associated with somatic mutation rates (Schuster-Bockler and Lehner 2012). On a much finer scale, the individual binding sites of a variety of DNA binding factors, including CTCF, appear to obstruct the lagging strand replication and DNA repair machinery and induce higher mutation rates in human and yeast (Reijns et al. 2015; Perera et al. 2016; Sabarinathan et al. 2016). However the mutational landscape associated with intermediate levels of chromatin organisation, such as chromatin loops, are not well studied.

During meiosis, recombination is initiated by double-strand breaks (DSBs) and occurs non-randomly across genomes; it is at its highest level at recombination hotspots (HSs) where the majority (60%) of recombination events take place (Myers et al. 2005; Coop et al. 2008). While it is known that recombination produces large structural variants, the effect of recombination on the emergence of single nucleotide variants is less clear - as is its relation to chromatin structure. There is evidence that recombination is mutagenic in yeast (Strathern et al. 1995; Hicks et al. 2010), and a recent study of 283 human trios has shown a correlation between the rate of recombination events in parental germ cell genomes and the rate of *de novo* SNVs in offspring genomes, suggesting a mutagenic effect of HSs (Besenbacher et al. 2016). However, the data supporting this were necessarily sparse, given the low *de novo* mutation rates in the normal human genome. Replication and recombination associated mechanisms are also hypothesised to lead to the formation of structural variants during mitosis, and may therefore contribute to structural variation in cancers (Carvalho and Lupski 2016). The influence of genomic features on structural rearrangements in cancer is relatively under-studied, but it seems that different cancer types follow different patterns. For some cancer types, such as breast cancer, structural somatic variants are enriched within early replicating, GC rich, transcribed regions of the genome (Drier et al. 2013), whereas the opposite trend was observed for cancers such as prostate and melanoma.

Here, we explore the unusual mutational landscape of LAPs, discovering significant depletions of SNVs at LAPs across multiple tumour types. Using matched replication timing and chromatin data, we find that these depletions can be explained by the mutational biases associated with LAPs. Paradoxically, we show that LAPs are also associated with increased rates of SVs in tumours and often overlap somatic DSB hotspots and meiotic recombination HSs. Like canonical common fragile sites, we find that LAPs are bound by components of DNA repair complexes such as BRCA1 and RAD51, and that BRCA-deficient tumours show different SNV rates at LAPs compared to wildtype tumours. LAPs which overlap recombination HSs are associated with increases in single nucleotide polymorphism (SNP) rates in extant human populations, suggesting that particular classes of LAPs act as reservoirs of sequence variation for evolution in the recent human lineage. We conclude that the unusual chromatin environment at LAPs influences the mutation rates observed and causes LAPs to be foci of evolutionary change.

## RESULTS

Previous work has demonstrated elevated SNV rates at CTCF binding sites within LAPs in a variety of cancers (Kaiser et al. 2016). This motivated us to investigate genome-wide somatic mutation rates around high confidence LAPs from the aggregated Hi-C datasets (see Methods) of Rao et al. (2014), using 13 recently released ICGC somatic variant datasets ascertaining both SNVs and SVs in 9 different tumour types (International Cancer Genome et al. 2010). The ICGC pan-cancer SNV rates show a dramatic drop within 50 Kb of LAPs (Figure 1A). This regional decrease in SNVs at LAPs is in stark contrast to the high mutation rate observed at the short 19bp binding CTCF-motifs located inside LAPs of 12.3 SNVs/Mb^-1^, or >3 times higher than the local background mutation rate. Plotting SNV rates at 20bp resolution, a peak of SNVs in the centre of the LAPs at CTCF binding sites becomes apparent (as seen in Kaiser et al. (2016)), as well as a periodic pattern of mutation reflecting nucleosome occupancy (Figure 1B) observed previously in breast tumour data (Morganella et al. 2016). Thus, CTCF binding sites within LAPs are prone to local somatic hypermutation in tumours, but, unexpectedly, these sites often reside within broader genomic regions with significantly reduced SNV rates.

**Figure 1:**
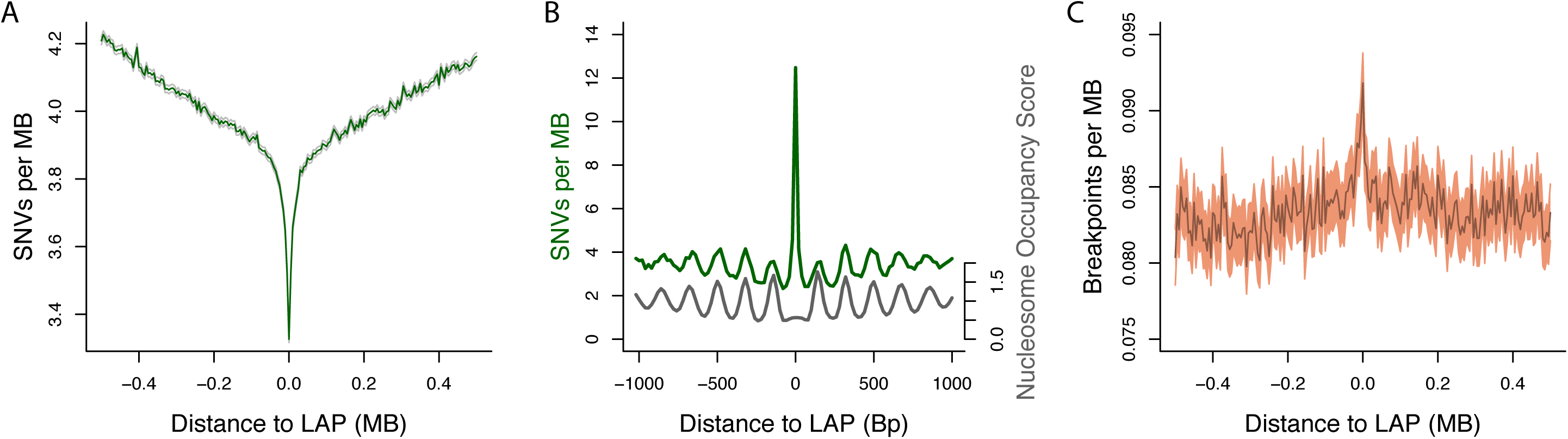
Loop anchor points are sites of genomic instability in cancer. (A) Average SNV rates per tumour within 500 Kb of all LAPs. (B) Finer scale average SNV rates per tumour within 1 Kb of LAPs, centred on the CTCF site within each LAP, with nucleosome occupancy based on MNase-seq data over the same intervals. (C) Average SV breakpoint rates per tumour within 500 Kb of all LAPs. 95% Confidence intervals, calculated assuming a Poisson distribution of mutation rates, are shown as shaded regions in (A) and (C).

### Chromatin loop anchors are hotspots of structural variation in tumours and recombination in human populations

In contrast to SNVs, the frequency of pan-cancer SV breakpoints shows a significant increase at LAPs, inverting the pattern seen for SNVs over the same range of flanking sequence (Figure 1C). This implies that LAPs are structurally fragile sites in cancer, and so we examined associations between LAPs and more direct measures of genomic instability. Lensing et al. (2016) identified genome-wide foci of endogenous DSBs *in vitro*, and these sites show a striking ~3.7-fold enrichment at LAPs (Figure 2A). A proportion of this enrichment may be attributable to the close proximity of LAPs to promoters and enhancers, which are known to suffer elevated DSB rates (Lensing et al. 2016). However, LAPs lacking any overlap with known promoters and enhancers show similarly elevated rates to those that do (Supplemental Figure S1). Consistent with this, LAPs are also enriched in predicted G-quadruplexes (G4s), a DNA secondary structure associated with regulatory regions and DSB formation in cancers (Hänsel-Hertsch et al. 2017) (Figure 2B). To our knowledge this is the first demonstration that LAPs are sites of inherent genomic instability, associated with an accumulation of SVs in their vicinity in tumour cells (Gaillard et al. 2015).

**Figure 2:**
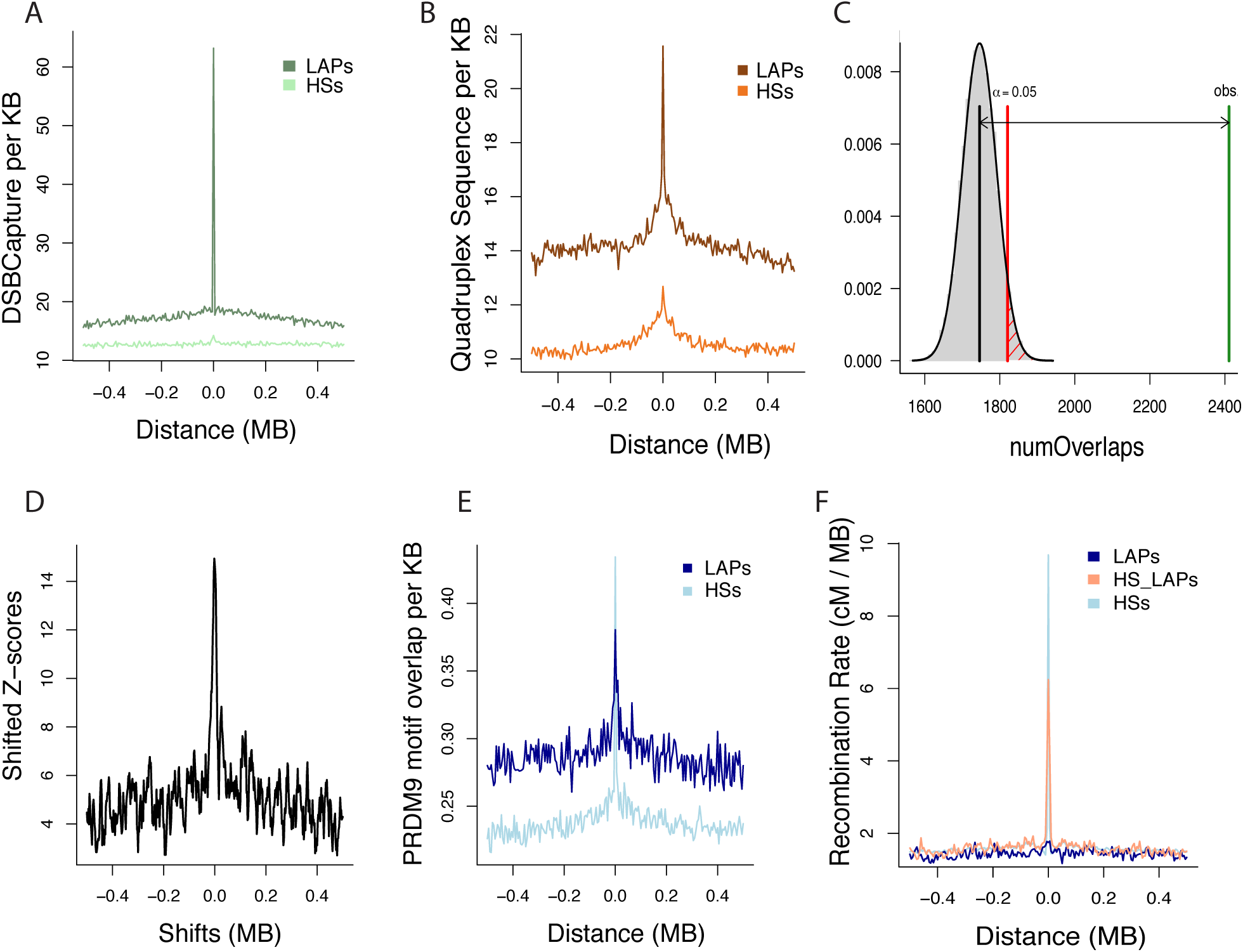
Loop anchor points are susceptible to somatic double strand breaks and meiotic recombination. (A) DNA double-strand break density within 500 Kb of LAPs and recombination HSs. (B) Predicted quadruplex structure forming sequence density within 500 Kb of LAPs. (C) Probability density plot showing the observed number (green line) *versus* expected distribution (grey histogram) of overlaps between LAPs and HSs, *p* < 10^-5^ based upon 100,000 circular permutations. The threshold for statistical significance is indicated by the red line. (D) Variation in circular permutation z-scores (y-axis) relative to shifts in the location of HSs (x-axis), suggesting that significance of overlaps between LAPs and HSs results from specific local (not broader regional) overlaps. (E) PRDM9 binding motif density, which is targeted by the recombination machinery, within 500 Kb of recombination HSs and LAPs. (F) Average recombination rates within 500 Kb of recombination HSs, the subset of LAPs overlapping HSs (HS_LAPs) and all LAPs.

Intriguingly, LAPs also show an unexpected genome-wide correspondence with germline recombination HSs, calculated from genotyping of extant human populations (The 1000 Genomes Project Consortium et al. 2015), such that 16% of LAPs overlap HSs (*p* < 10^-5^; Figure 2C). These overlaps are notably precise, so that the association between LAPs and HSs drops when the two sets of regions are shifted with respect to each other by less than 50 Kb (Figure 2D). Thus, this correspondence is not simply attributable to the enrichment of both sets of features within certain broader neighbourhoods such as replication timing domains or nuclear compartments. Recombination HSs are known to often contain the motif bound by PRDM9, a critical component of the recombination machinery (Myers et al. 2010; Pratto et al. 2014), and, using stringent search criteria (see Methods), we find this motif in 17% of HSs. Similarly, we find that 13% of LAPs also contain at least one PRDM9 core motif, which is an enrichment of ~33% compared to the median number of motifs per 5 Kb bin in LAP flanking regions (Figure 2E). For the 16% of LAPs directly overlapping HSs (HS-LAPs) there is a notable increase in the recombination rate measured at those LAPs, but, beyond these HS-LAPs, there is no evidence for increased levels of recombination at LAPs in the germline (Figure 2F). This is consistent with dual roles for a subset of LAPs, both as units of chromatin organisation and as hotspots of structural variation.

Although PRDM9 is normally expressed exclusively in testis, it is also expressed in a variety of cancer cell lines and samples, and has been proposed as a cancer biomarker (Feichtinger et al. 2012). We observe modestly increased SNV rates at recombination HSs in cancer, but do not find any pan-cancer increase in SV breakpoints around HSs (Supplemental Figure S2), which might be expected if meiotic recombination complexes were activated in the tumours examined here. In addition, the histone modification H3K4me3, which is deposited by PRDM9 at DSBs, is not observed at recombination HSs in the HepG2 cell line (Supplemental Figure S3) nor in MCF-7 (data not shown). In contrast, H3K4me3 increases around LAPs – possibly as a result of PRDM9 recruitment to the PRDM9 motifs enriched at LAPs – or, alternatively, because H3K4me3 is a mark of active promoters enriched at chromatin boundaries (Moore et al. 2015) (Supplemental Figure S3). We cannot, however, exclude the possibility that PRDM9 is active in at least a subset of the tumours under investigation, and is responsible for the increase in SV rates at LAPs.

A substantial fraction of LAPs (47% of those studied here) constitute regulatory domain boundaries (Rao et al. 2014), while an even higher proportion, 69%, overlap DSB-foci (Lensing et al. 2016), and 16% overlap recombination HSs (Table 1). However, genome-wide, these three categories of LAPs appear to be largely independent, as the extent of overlap between categories was remarkably similar to the expected rate assuming independent distributions across the genome. For example, LAPs that appear as domain boundaries were as likely to overlap recombination HSs as LAPs that do not act as boundaries (Table 1). Furthermore, there was no enrichment of GO terms associated with the genes neighbouring HS-LAPs *versus* a background set of those found at all LAPs, i.e. HS-LAPs are not found near specific functional categories of genes.

**Table 1:** Overlap of LAPs with domain boundaries, recombination HSs, and DSB-prone ‘Breakome’ regions. A) The number of features and their genomic span. B) Overlap statistics. Columns show the feature intersection, the numbers and fractions (as a proportion of the total dataset of 14,737 LAPs) of unique LAPs in each intersection, and the expected fraction of overlap under independence of features.

| A) | Feature | Feature Count | Genomic Span (Mb) |
| --- | --- | --- | --- |
|  | LAPs | 14,737 | 73.69 |
|  | Boundaries | 40,615 | 250.89 |
|  | HSs | 32,984 | 181.79 |
|  | Breakome | 84,946 | 34.69 |

| B) | Feature Intersection | Number of LAPs | Fraction of LAPs | Expected |
| --- | --- | --- | --- | --- |
| | LAPs ∩ Boundaries | 6,960 | 0.47 | - |
| | LAPs ∩ HSs | 2,385 | 0.16 | - |
| | LAPs ∩ Breakome | 10,133 | 0.69 | - |
| | LAPs ∩ HSs ∩ Breakome | 1,709 | 0.12 | 0.11 |
| | LAPs ∩ HSs ∩ Boundaries | 1,172 | 0.08 | 0.08 |
| | LAPs ∩ HSs ∩ Boundaries ∩ Breakome | 836 | 0.06 | 0.05 |

Although the three categories of LAPs (those annotated as domain boundaries, DSB regions and recombination HSs) occur to a large degree at distinct locations, they are associated with similar mutational landscapes in tumours. All three categories show the distinctive dip in SNV rates at LAPs (Figure 3A), while cancer mutation rates somewhat increase around recombination HSs, by ~3.5% compared to the median rate within the flanking regions. As expected, recombination HSs are associated with a pronounced increase in SNPs in the 1KG dataset, and LAPs that overlap HSs are also enriched for segregating variants in the 1KG dataset, by ~7% compared to the flanking regions (Figure 3B). Accordingly, population genetic processes that increase variation at HSs - such as selective sweeps and reductions in background selection (Charlesworth 2009) - also appear to have impacted germline variation at HS-LAPs. However, germline *de novo* SNV rates at LAPs are not reduced as they are in cancer and, similarly, we do not observe an increase in SVs near HSs (Supplemental Figure S2), suggesting fundamentally different influences on germline mutation rates *versus* cancer associated somatic mutation rates at these sites.

**Figure 3:**
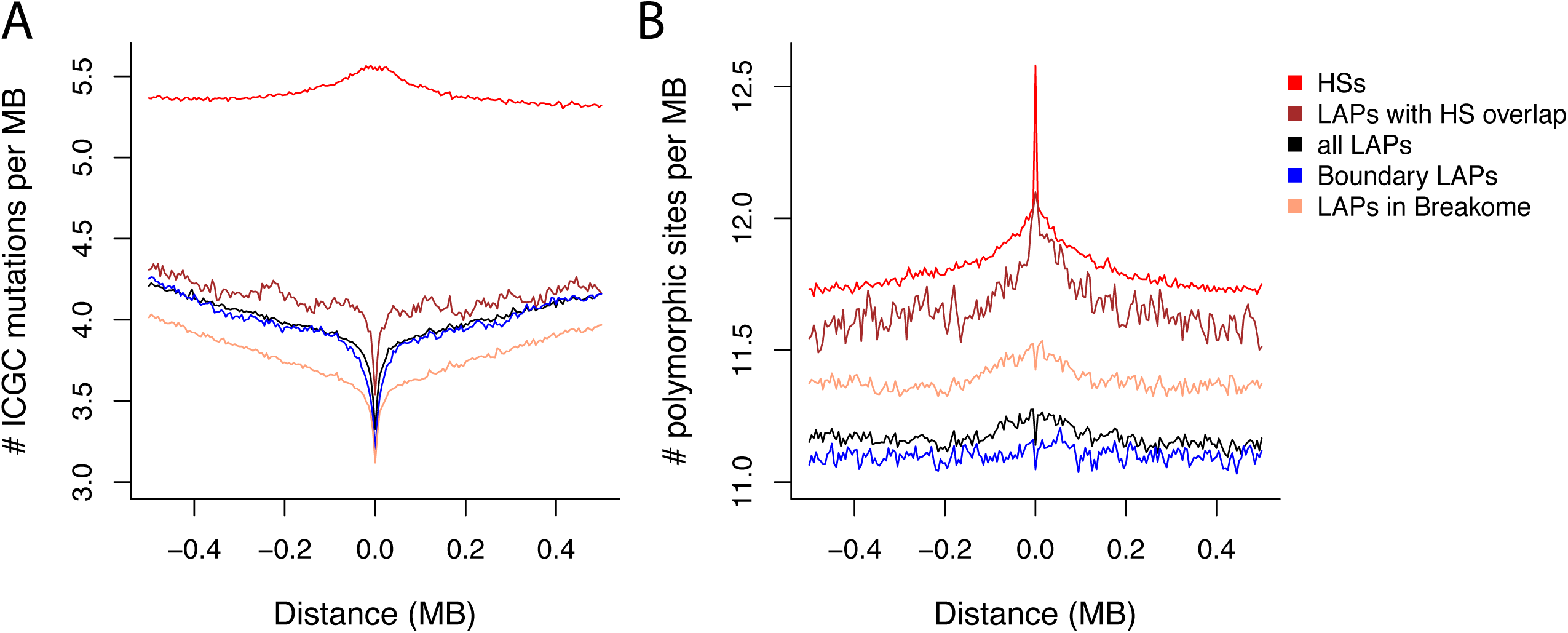
The impact of loop anchor points on tumour and population variation. (A) Average SNV somatic mutations based upon ICGC pan-cancer tumour sequencing data and (B) the average number of polymorphic segregating sites within 500 Kb of LAPs and HSs. LAPs are subdivided into those that overlap recombination HSs, domain boundaries or DSB regions (LAPs in Breakome).

### Chromatin features influence increased mutation rates at LAPs

LAPs are relatively GC-rich, enriched for histone modifications associated with active transcription (H3K27ac, H3K27me3, H3K36me3, H3K4me3, and DNase sensitive open chromatin) and depleted for the repressive mark H3K9me3; they also tend to be found in relatively early replicating regions of the genome (Supplemental Figure S3). Recombination HSs also show locally increased GC content, but otherwise generally possess a contrasting set of features, consistent with their presence in later replicating regions (Supplemental Figure S3). Accordingly, LAPs tend to be enriched for genes and actively transcribed regions (Rao et al. 2014), whereas HSs are located, on average, further away from genic sequence. This raises the possibility that the unusual mutational properties of LAPs may be explained by their distinctive chromatin and sequence features.

We used random forest regression models to assess the extent to which mammary epithelium derived chromatin features (from the MCF-7 and MCF10A cell lines) and a variety of other features were associated with mutation rates in a large ICGC breast tumour dataset (BRCA-EU). Specifically, we constructed models of mutation rates observed within all 5 Kb windows from the 500 Kb regions flanking all mammary epithelium LAPs (derived from HMEC cell line Hi-C data) plus the 5 Kb LAP regions themselves (Methods). Similar models have previously shown high predictive accuracy in modelling aspects of nuclear organisation and provide variable importance estimates that are robust to the inter-correlated nature of chromatin feature input variables (Moore et al. 2015). In our model, by far the most important predictor of the BRCA-EU SNV rate was replication timing, with reduced levels of mutation observed in early replicating regions (Figure 4), consistent with other studies of breast cancer mutation patterns (Morganella et al. 2016). Notably, the allocation of a genomic region to a LAP had little impact on the SNV rate, i.e. changes in mutation rates were mostly explained by the genomic features at LAPs, rather than by LAP presence. The correlation coefficient between observed and predicted SNV rates from the random forest model (*r* = 0.28; *p*-value < 10^-15^) suggests a significant influence of the features included but, overall, a moderate level of predictive accuracy. Modelling was less successful in predicting BRCA-EU SV breakpoint rates (*r* = 0.09 between observed and predicted values) but also indicated a significant association with chromatin and sequence features (*p*-value < 10^-15^), most notably BRCA1 (Figure 4B). However, the effects of most features on mutation rates are strikingly inverted for SNV and SV rates, such that the variables most strongly correlated with elevated SV rates (DNaseHS, replication timing, G-quadruplex content, GC content) are associated with decreased SNV rates (Figure 4C). We conclude that similar chromatin and sequence features have significant, but largely opposing, effects upon SNV and SV rates at LAPs.

**Figure 4:**
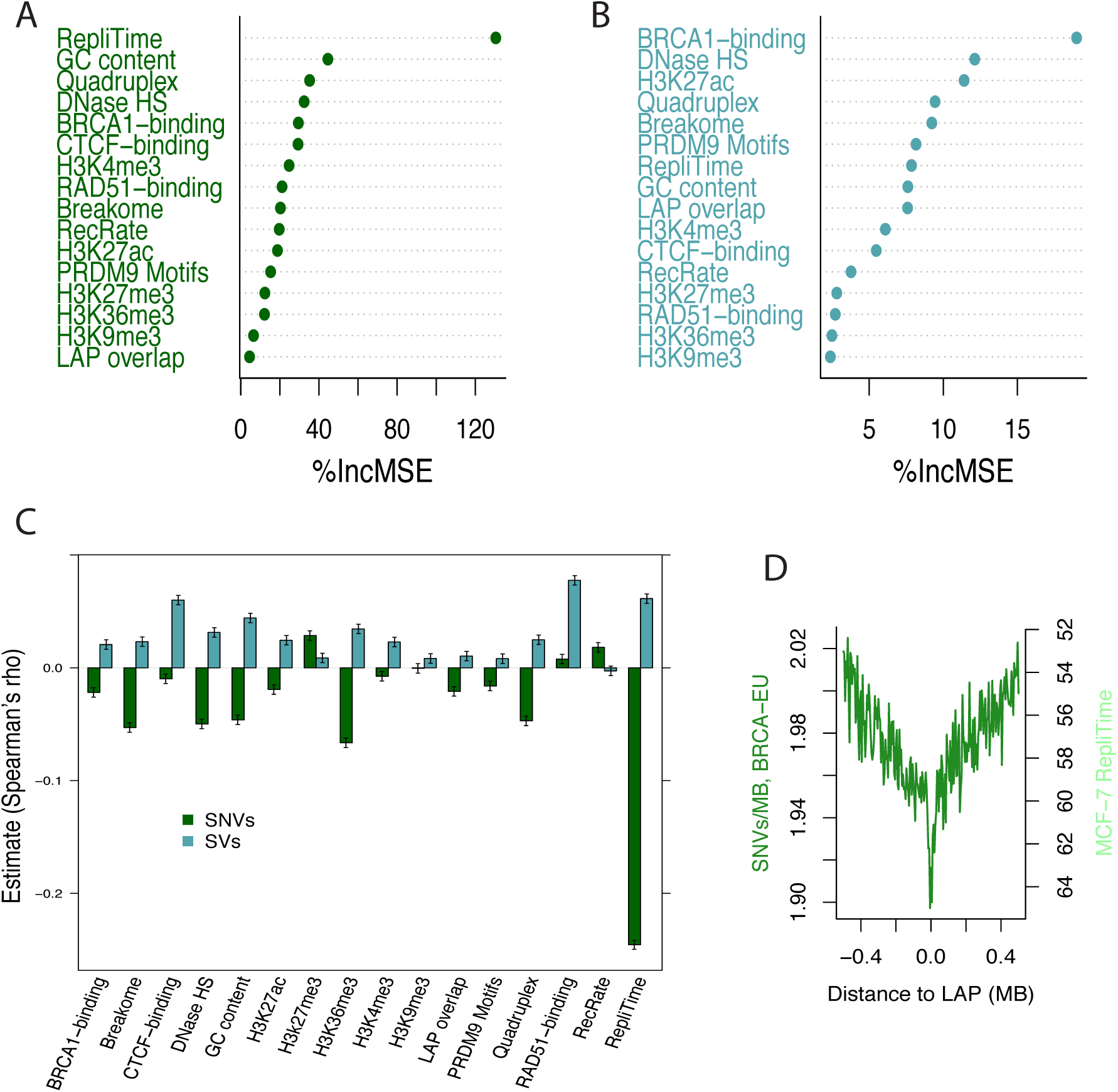
Genomic and epigenomic features influencing SNV and SV rates at loop anchor points in breast tumours. (A, B) Variable importance, measured as percentage increase in mean squared error (%IncMSE) when each predictor variable is removed from the model, predicting SNV rates (A) or SV breakpoint rates (B), respectively. (C) Spearman’s rank correlation coefficient estimates between mutation rates and predictor variables of the random forest models, for SNV rates (green) and SV breakpoint rates (blue); 95% confidence intervals of the estimates are indicated. (D) SNV rates in BRCA-EU breast tumours within 500 Kb of LAPs in the MCF-7 breast tumour cell line relative to replication timing (average MCF-7 ENCODE Repli-seq signal); higher Repli-seq values indicate earlier replication time.

The modest overall predictive power of the SNV and SV random forest models undoubtedly reflects the necessarily incomplete nature of the chromatin and sequence features included as input, omitting many currently unavailable variables. Additional limitations are expected to result from the current sample sizes of variant datasets, limiting the resolution of the modelling to 5 Kb regions that are much larger than many of the chromatin and sequence features. In support of this, smaller sub-regions within HMEC LAPs annotated as insulator, promoter or enhancer sites in HMEC cells (see Methods) differ significantly in their mutation rates and replication timing. Insulator sites show later average replication timing in MCF-7 cells than regions within LAPs classified as promoters or enhancers (Supplemental Figure S4). Concordant with this, the per-Mb SNV rates were 2.23, 1.93 and 1.92 for insulator, promoter and enhancer regions, respectively (Table S1; relative proportion tests *p* < 10^-7^ for the comparisons “insulator *versus* promoter” and “insulator *versus* enhancer”). The equivalent SV breakpoint rates per Mb suggest an increase in the relatively early replicating promoter regions (Table S1), again, consistent with the broad trends seen in the models (Figure 4).

Our data benefits from aggregate analyses of mutation rates across cancer types and LAPs, allowing us to highlight common patterns of mutation. Supplemental Figure S5 shows SNV and SV rates around LAPs for all nine cancer types in this study separately. SNV rates are consistently reduced near LAPs in several tumour types, but there are exceptions, such as the malignant lymphoma (MALY-DE) dataset, which does not show a pronounced dip in mutation rates near LAPs though it includes a large number of SNVs (Table S2). Beyond differences in SNV dataset sample sizes, this variability among tumour types may reflect the limitations of the current Hi-C data, which may be poorly matched to the cells in certain cancer samples. In addition, SVs are, on average, ~100-fold less frequent than SNVs (Supplemental Table S2), and stratifying SVs by tumour type reduces the power to detect any patterns on a per-tumour basis.

### BRCA1/2 deficient breast tumours suffer higher mutation loads at LAPs

BRCA1 and BRCA2 are well-characterised tumour suppressor genes involved in DSB repair by homologous recombination (Deng and Wang 2003; Lord and Ashworth 2012; Walsh 2015). BRCA1 is often recruited to sites of active transcription, which are prone to DNA damage during the formation of transcriptional R-loops (Hatchi et al. 2015). We found a strong enrichment of BRCA1 at HMEC LAPs (but not HSs) in MCF10A, a normal breast epithelial cell line, as well as increased RAD51 binding (which mediates BRCA2 binding) around LAPs in the MCF-7 cell line (Figure 5). BRCA1-binding at LAPs in MCF10A cells was a relatively influential predictor in the random forest SV model (above), such that higher levels of BRCA1 binding was associated with higher SV rates (Figure 4). To our knowledge, this is the first observation of BRCA1/2 association with LAPs, and with mutation rates at LAPs. To further investigate the importance of BRCA1/BRCA2 binding at LAPs, we exploited a recent classification of BRCA-EU breast cancer tumours as BRCA1/BRCA2 deficient (Davies et al. 2017).

**Figure 5:**
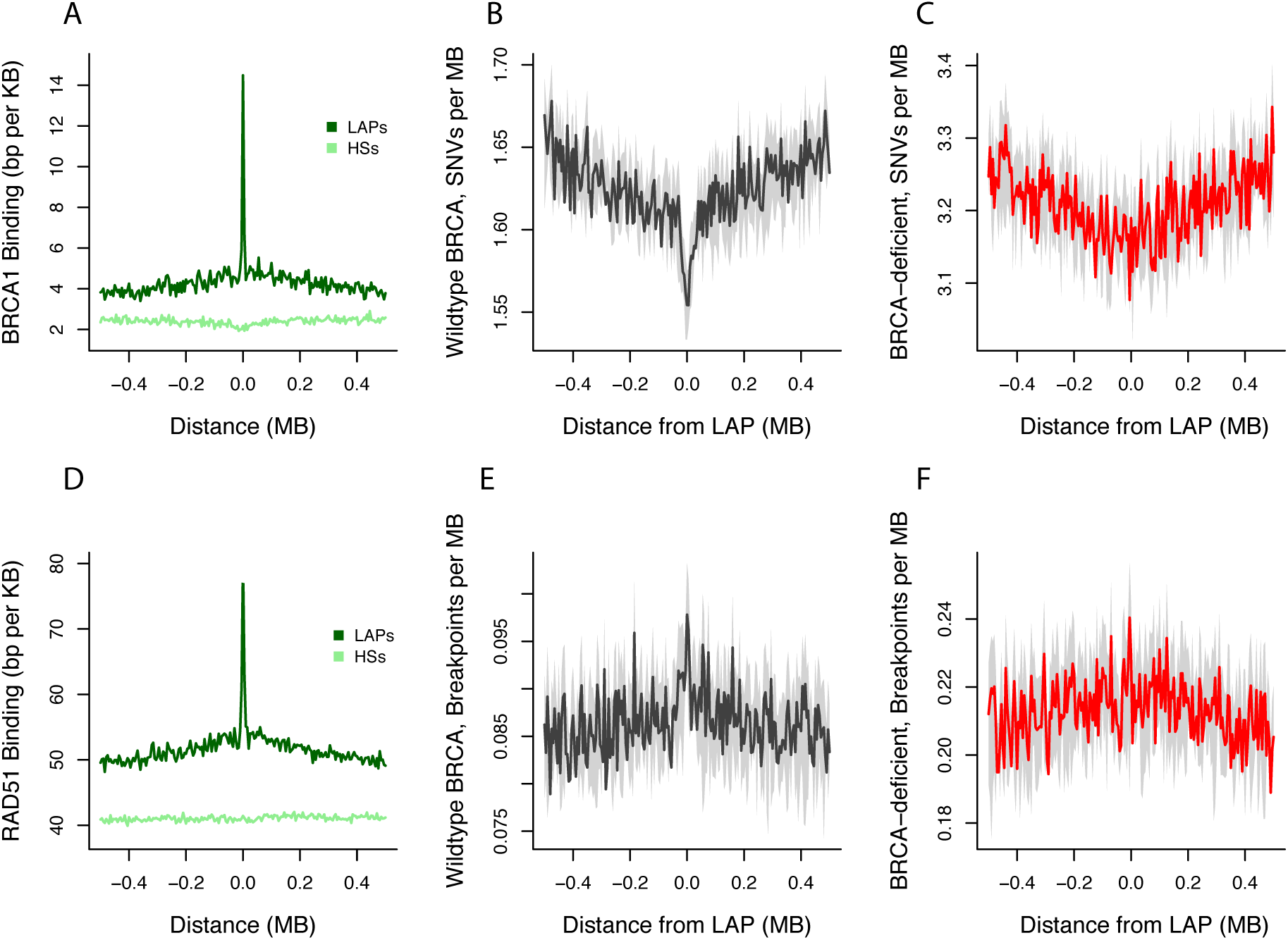
BRCA-deficiency influences mutation rates at LAPs. (A) Average density of BRCA1 ChIP-seq binding peaks for MCF-7 cells and (D) RAD51 ChIP-seq binding peaks for MCF-10A genome-wide for LAPs and recombination HSs. Breast tumour SNV rates within 500 Kb of normal breast epithelial cells (HMEC) LAPs for BRCA wildtype tumours (B) and BRCA deficient tumours (C). Breast tumour SV breakpoint rates within 500 Kb of HMEC LAPs for BRCA wildtype (E) and BRCA deficient (F) tumours.

BRCA1/BRCA2 deficient samples have higher genome-wide SNV rates, but a less pronounced dip in mutation rates around HMEC LAPs compared with BRCA wild-type tumours (Figure 5). In the BRCA deficient samples, 1.31% of all somatic SNVs were found at HMEC LAPs (16,154 out of 1,229,533 mutations across 124 patients), whereas the corresponding fraction was significantly smaller, 1.24%, in tumours with intact BRCA (27,856 out of 2,250,119 mutations across 436 patients) (two-sided Fisher’s exact test; *p* < 10^-8^). Furthermore, SV breakpoint rates more than doubled in BRCA deficient samples compared to wild-type tumours genome-wide (on average, there were 206 and 527 breakpoints per BRCA deficient and wild-type sample, respectively), but relatively fewer breakpoints occurred at LAPs in the BRCA-deficient samples (two-sided Fisher’s exact test; *p* < 0.03) (Figure 5E and 5F). We conclude that deficiencies in BRCA1/BRCA2 are associated with an increase of SNVs at LAPs (reflected in a less pronounced dip in SNV rate) and a general increase in SV generation that is not confined to LAPs.

These (BRCA-EU) breast tumours show a strong C>G and C>T signature at Tp<u>Cp</u>N sites which is less pronounced for BRCA1-BRCA2 deficient tumours (Supplemental Figure S6A), as expected since the classification of these tumours was largely based upon SNV mutation signatures (Davies et al. 2017). While most of the variation in mutational input was explained by just two signatures that separate the wild-type and deficient tumours, signatures 3 and 4 separated mutations that occurred at LAPs and control sites (Supplemental Figure S6). That is, we observe a shift in the types of mutations that occur at LAPs compared with control regions in both the deficient and wild-type tumours.

### Genes within recurrently disrupted chromatin loops are enriched for functions in the cell cycle

Recent literature has documented tumours showing oncogene upregulation as a result of disrupted CTCF binding sites and chromatin loops in a variety of cancers (Kaiser and Semple 2017). Using breakpoint data from all 1,672 ICGC donors and the set of 14,737 high confidence LAPs, we find that genes within the top 5% most disrupted chromatin loops (possessing 5 or more SV breakpoints in their LAPs) are indeed enriched for functional annotation terms associated with proliferation and the G1/S cell cycle transition (Table 2) (Bertoli et al. 2013). The enrichments of such putatively cancer-associated terms are often only marginally significant given current sample sizes, but are broadly consistent with the previously reported oncogenic disruptions (Kaiser and Semple 2017). Subdividing the dataset into more refined categories of breakpoints and LAPs (e.g. LAPs affected by deletions or duplications only) does not yield a stronger enrichment of terms (data not shown). Thus, it is possible that frequent disruptions of chromatin loops and domain boundaries in proximity to oncogenes in tumours are driven by the unusual mutational biases at LAPs.

**Table 2:**
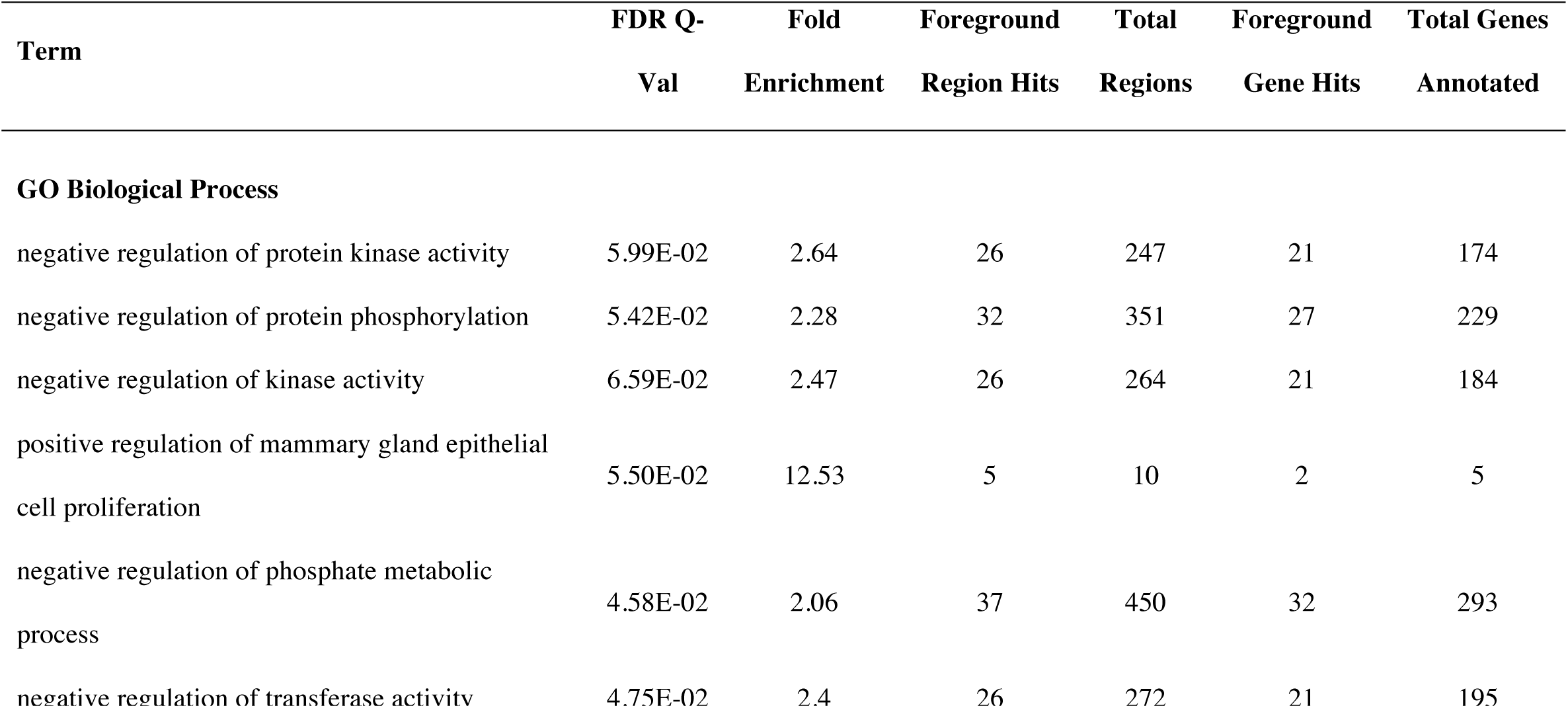
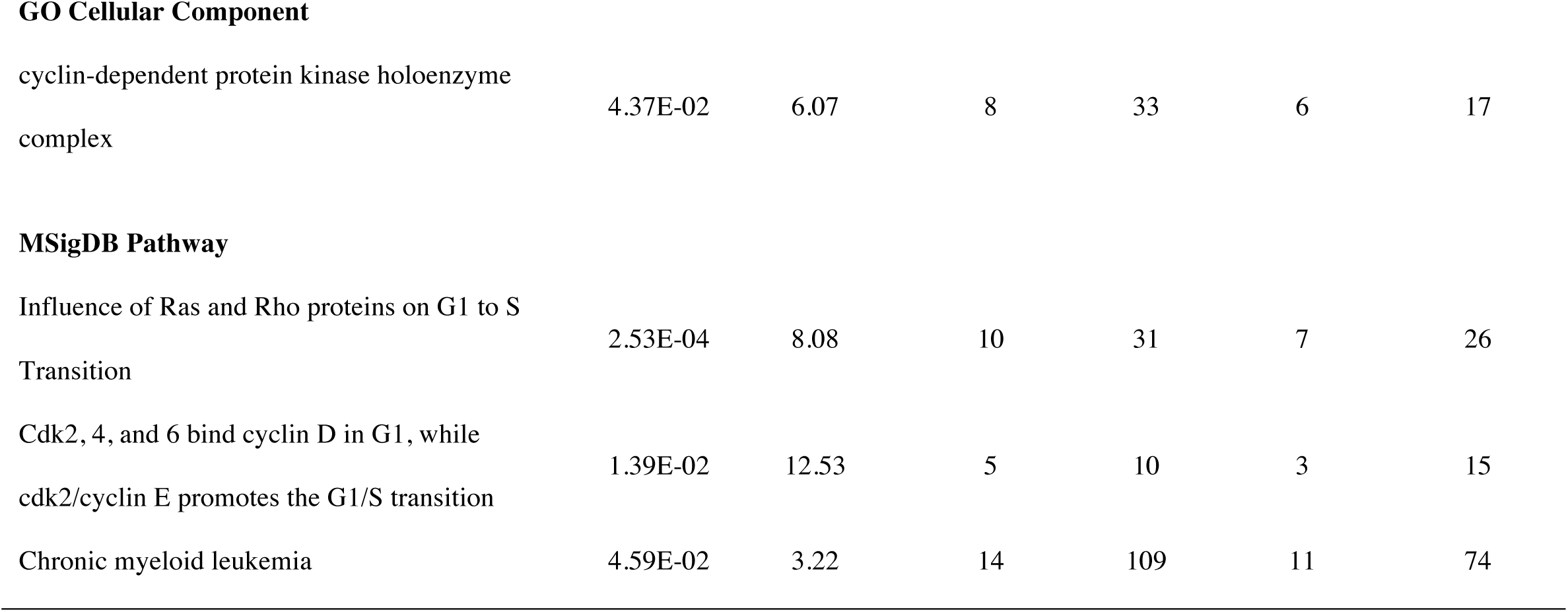
Functional annotation enrichments at recurrently disrupted LAPs. Enrichment of genes within the foreground set of 398 genomic regions (chromatin loops recurrently affected by structural variant breakpoints), relative to the background set (9,973 loops defined by all 14,737 LAPs) for GO and pathway terms. All regions were associated with genes using the GREAT tool (see Methods).

| Term | FDR Q-Val | Fold Enrichment | Foreground Region Hits | Total Regions | Foreground Gene Hits | Total Genes Annotated |
| --- | --- | --- | --- | --- | --- | --- |
| **GO Biological Process** |  |  |  |  |  |  |
| negative regulation of protein kinase activity | 5.99E-02 | 2.64 | 26 | 247 | 21 | 174 |
| negative regulation of protein phosphorylation | 5.42E-02 | 2.28 | 32 | 351 | 27 | 229 |
| negative regulation of kinase activity | 6.59E-02 | 2.47 | 26 | 264 | 21 | 184 |
| positive regulation of mammary gland epithelial cell proliferation | 5.50E-02 | 12.53 | 5 | 10 | 2 | 5 |
| negative regulation of phosphate metabolic process | 4.58E-02 | 2.06 | 37 | 450 | 32 | 293 |
| negative regulation of transferase activity | 4.75E-02 | 2.4 | 26 | 272 | 21 | 195 |
| **GO Cellular Component** |  |  |  |  |  |  |
| cyclin-dependent protein kinase holoenzyme complex | 4.37E-02 | 6.07 | 8 | 33 | 6 | 17 |
| **MSigDB Pathway** |  |  |  |  |  |  |
| Influence of Ras and Rho proteins on G1 to S Transition | 2.53E-04 | 8.08 | 10 | 31 | 7 | 26 |
| Cdk2, 4, and 6 bind cyclin D in G1, while cdk2/cyclin E promotes the G1/S transition | 1.39E-02 | 12.53 | 5 | 10 | 3 | 15 |
| Chronic myeloid leukemia | 4.59E-02 | 3.22 | 14 | 109 | 11 | 74 |

In the breast cancer dataset, we do observe an unexpected excess of overlap between recurrently (> 2x) disrupted HMEC loops and GWAS regions associated with breast cancer: There were 40 such overlaps, whereas only 18.1 overlaps were, on average, observed in 5,000 circular permutations (*p* = 0.0002). This excess in overlap is notably larger than that observed for the background set of all HMEC loops and GWAS hits (225 observed overlaps and a mean of 143.6 expected overlaps, based on 5000 permutations; *p* = 0.001), suggesting a possible causal relationship between LAP disruption and the breast cancer phenotype. However, in common with previous studies (Kaiser and Semple 2017), it is unclear to what extent this relationship is driven by the mutational biases that we have demonstrated at LAPs or selective processes in tumours.

## DISCUSSION

LAPs and recombination HSs are two seemingly unrelated features of the genome - one involved in chromatin organisation and the other in recombination during meiosis - but both classes of sites emerge as hotspots for DSBs. We have shown that LAPs and HSs often occur in the same genomic locations, which suggests that the same genomic regions that migrate to the chromosomal axis during meiosis, ultimately forming the points of breakage for DSB initiation (Baudat et al. 2013), are also involved in chromatin organisation in the interphase nucleus of somatic cells. Intriguingly, cohesin, which associates with CTCF at LAPs (Ong and Corces 2014; Tang et al. 2015), is also enriched at the meiotic loop axis and plays a diverse role in chromosome pairing in both mitosis and meiosis (McNicoll et al. 2013). A variety of factors, many related to chromatin structure, affect the propensity of LAPs to harbour SV breakpoints. PRDM9 also appears to be active in at least a subset of cancer cells (Feichtinger et al. 2012) and may contribute to DSB formation at LAPs, suggesting another possible link between LAPs and HSs. The association of LAPs with DSB formation appears to be at least partly attributable to the enrichment of active promoters and enhancers at LAPs, which is consistent with reports that promoters are inherently prone to DSBs, both in somatic cells (Lensing et al. 2016) and meiotic cells that lack PRDM9 (Brick et al. 2012). However, we also observe an excess of DSBs at LAPs showing no overlap with promoter or enhancer states, and our modelling suggests that the presence of LAPs influences SV rates independently of features associated with these states. LAPs are also unusual with respect to replication timing, with LAPs replicating, on average, earlier than their surrounding regions – consistent with the dual roles of cohesin in stabilising chromatin loops and also initiating replication (Guillou et al. 2010). Accordingly, chromatin looping may, to some extent, directly result from the initiation of replication, or, conversely, determine its starting position in the next cell cycle (Courbet et al. 2008). Given the strong association between LAPs and DSB-prone regions, disruption of replication near such origins may be one way in which genome instability is introduced in cancer (Losada 2014). Notably, regions stably bound by DNA binding proteins such as CTCF seem to suffer high mutational loads due to replication errors (Reijns et al. 2015).

LAPs are foci of DSB breakpoints and may provide the raw material for cancer evolution via structural variation, dependent on other factors, such as deficiencies in DNA repair pathways. The resulting SVs may have be subject to selection in cancer, though the majority are likely to be ‘passenger’ variants that drift toward fixation with little phenotypic consequences in their tissue of origin. Accordingly, we observe a strong enrichment of somatic DSB formation at LAPs in the NHEK cell line, more modest elevations in SV breakpoints around LAPs in cancers, and limited evidence that the genes affected are those that experience selection in cancers. At the human population level, our results suggest that chromatin loops are predominantly inherited as a genetic unit, with recombination often confined to LAPs, and therefore tending to preserve regulatory haplotypes. Consistent with our results, recent studies have shown that LD blocks are enriched within topologically associating domains, i.e. recombination between enhancers and their target genes is reduced within domains (Liu et al. 2016). Indeed, HSs themselves may primarily be a by-product of particular chromatin environments and other functional constraints, such as a lack of active transcription, which may interfere with the recombination process (McVicker and Green 2010).

We have used aggregate analysis across loop anchor points and cancer types, to show that a mutational bias towards breakage of chromatin loop anchors exists; notably, it is more prominent in some cancer types than others and presumably depends on the general genome instability of the tumour type. The unusual somatic DNA breakage patterns near LAPs are likely to contribute to cancer evolution, reflected in higher SV breakpoint levels, and allowing novel promoter-enhancer interactions. The increase in breakage near LAPs is influenced by their specific chromatin environment and replication timing, DNA folding and accessibility to the repair machinery. Similar influences may underlie the surprising association of LAPs with meiotic recombination events.

## METHODS

### Datasets

Chromatin loops for cell lines representing all human germ layers (GM12878, HeLa, HMEC, HUVEC, IMR90, K562 and NHEK) were derived from unusually high resolution *in situ* Hi-C data, defining LAPs at a resolution of 1-5 Kb (GSE63525; (Rao et al. 2014)). These loops are often conserved between cell lines, such that 55-75% of the loops detected in any given cell line were also found in the most deeply sequenced cell line (GM12878), and around 50% appear to be conserved across mammalian species (Rao et al. 2014). The majority of loops are also associated with convergently orientated CTCF binding motifs at the putative LAPs, consistent with the known roles of CTCF in loop formation (Rao et al. 2014). On average, 17% of LAPs were only observed in one tissue (Supplemental Figure S7), with more deeply sequenced cell lines consistently resulting in more LAPs being called. From this dataset, we created a merged dataset of 14,737 LAPs, centred around their associated (and convergently orientated) CTCF motifs (Rao et al. 2014), which represents the union of high confidence LAPs across all cell lines. This merged dataset was used for all analyses except for the breast cancer specific analysis, where we used only LAPs derived from the HMEC (human mammary epithelial cells) Hi-C data, and the DSB analysis where tissue-matched NHEK (normal human epidermal keratinocyte) LAPs were used.

Recombination hotspot locations had been identified in the Phase II HapMap dataset (release 21) (McVean et al. 2004; Winckler et al. 2005); recombination rates and SNP data were derived from Phase 3 of the 1000 Genomes project (The 1000 Genomes Project Consortium et al. 2015). For the mutation rate analysis, we created 500 Kb windows around the midpoints of LAPs and HSs, and omitted regions containing ENCODE blacklisted genomic regions (http://hgdownload.cse.ucsc.edu/goldenPath/hg19/encodeDCC/wgEncodeMapability), resulting in 11,085 union LAPs, 6,214 HMEC LAPs, and 18,914 recombination HSs plus their respective flanking regions. Median distances between the centre points of adjacent features were 106,331 bp for union LAPs, 125,000 bp for HMEC LAPs and 55,500 bp for HSs, respectively. We divided each 500 Kb region around a LAP into 5 Kb non-overlapping windows; measures of mutation rates and overlap with other genomic features were calculated for each 5 Kb bin.

ICGC (https://dcc.icgc.org/) datasets with whole genome sequence derived calls for both SNV and SVs (not under publication moratorium at the time of analysis) were included: BOCA-FR (bone cancer), BRCA-EU (breast cancer), CLLE-ES (chronic lymphocytic leukemia), LIRI-JP (liver cancer), MALY-DE (malignant lymphoma), MELA-AU (skin cancer), OV-AU (ovarian cancer), PACA-AU (pancreatic cancer), PAEN-AU (pancreatic cancer), PAEN-IT (pancreatic cancer), EOPC-DE (early onset prostate cancer), PRAD-CA (prostate cancer), PRAD-UK (prostate cancer). The combined analysis included a total of 32,105,808 SNVs and 368,480 structural variants (Table S2). We included all categories of structural variants that were listed in the ICGC files (i.e. insertions, deletions, inversions etc.), recording all breakpoint positions based on the coordinates of the SVs (i.e. a single SV has two breakpoint positions). SNV and breakpoint rates were intersected with the genomic coordinates of LAPs and HSs. Confidence intervals (as in Figure 1) were calculated based on the assumption that breakpoint and SNV rates are random processes and follow the Poisson distribution.

Germline *de novo* mutation rates were reported for a whole genome sequencing study of 283 Icelandic trios (Besenbacher et al. 2016). A range of genomic features were generated by the ENCODE consortium (Encode Project Consortium 2012), including average replication timing for each 5 Kb genomic window, which was calculated based on the Repli-seq wavelet-smoothed signal for MCF-7 (Breast cancer) and HepG2 (Liver cancer) cell lines; open chromatin sites (DNAse hypersensitivity), CTCF-binding in MCF-7; RAD51-binding in MCF-7; Broad chromHMM tracks for chromatin state segmentation of HMEC; histone modifications (H3K27ac, H3K09me3, H3K4me3, H3K36me3, and H3K27me3) in MCF-7 and HepG2; nucleosome occupancy scores for GM12878. Genomic features based on ChIP-seq data were represented as genomic segments (peaks called) in the ENCODE distributed files; the overlap (in basepairs) between these features and genomic windows around LAPs was calculated using bedtools (Quinlan and Hall 2010). GC-content for each 5 Kb window was also calculated using bedtools (Quinlan and Hall 2010). Sites predicted to adopt quadruplex conformations were generated by Kudlicki (2016) (http://moment.utmb.edu/allquads/). DSBs were detected using the DSBCapture protocol in the NHEK cell line (GEO database accession: GSE78172) (Lensing et al. 2016). ChIP-seq data for BRCA1 binding in MCF-10A cells were generated by Gardini et al. (2014) and MACS2 (Zhang et al. 2008) was used to call peaks in BRCA1-binding using default parameters.

### Modelling and analysis

Random Forest regression models were constructed using the R package randomForest (Liaw and Wiener 2002). To construct a model with 200 trees, we extracted genomic regions within 500KB of a HMEC LAP, merging overlapping regions, i.e. counting each unique genomic region once. Response variables were the number of BRCA-EU SNVs or SVs per 5-KB window, for a total of 239,141 windows. Predictor variables were replication timing in MCF-7; GC content; quadruplex overlap; HapMap recombination rate; DSB regions in NHEK cells; MCF-7 DNAse hypersensitivity; BRCA1-binding in MCF10A; RAD51-binding in MCF-7; CTCF-binding in MCF-7; PRDM9 motif coverage; overlap with peaks of H3K4me3, H3K27ac, H3K36me3, H3K27me3, H3K9me3 for MCF-7 and MCF-10A cell lines; HMEC LAP presence.

BRCA-EU Patient IDs with a predicted deficiency in BRCA1/BRCA2 were extracted from Davies *et al.* (Davies et al. 2017); the HRDetect probabilistic cut-off of 0.7 was used, resulting in 124 patients classified as BRCA-deficient, and 436 as BRCA wildtype (Davies et al. 2017). The R package SomaticSignatures (Gehring et al. 2015) was used to calculate mutational signatures at LAPs and control sites for breast cancer samples classified as BRCA normal and BRCA deficient. The number of signatures that best explain the variation in mutational profiles was calculated using the “assessNumberSignatures” function. Mutational signatures were compared between cancer SNVs that occurred at HMEC LAPs and 5 Kb control windows placed 1 Mb downstream from the LAPs. Circular permutation within the R package “RegioneR” (Gel et al. 2016) was used to assess the significance of genome-wide overlap between LAPs and recombination HS, using 100,000 permutations. The FIMO algorithm (Grant et al. 2011) from the MEME package (Bailey et al. 2009) was used to scan the genome for occurrences of the 13-bp PRDM9-motif CCTCCCTNNCCAC, using default parameters; this resulted in 51,107 motif locations being identified with a motif match p-value < 1.3e-06. Functional annotation enrichment analysis was carried out for regions of interest using the GREAT tool to calculate FDR corrected hypergeometric q-values for the default selection of annotation ontologies (McLean et al. 2010).

Breast cancer associated SNPs were obtained from the GWAS catalogue (MacArthur et al. 2017) (2017-05-29 release), and their coordinates were extended by 5 Kb (to account for LD tagging of nearby causal SNPs) according to the average span of LD blocks in 1000 Genome Project data for European populations (Rosenfeld et al. 2012). The resulting GWAS SNP-containing segments were merged using bedtools (Quinlan and Hall 2010) to create a non-redundant set of GWAS regions; circular permutations were carried out in R to test for an excess of overlap between the GWAS regions and chromatin loops in the HMEC cell line.

## DATA ACCESS

Data sharing not applicable: all data used in this study is publicly available, from sources indicated in the manuscript.

## ACKNOWLEDGMENTS

This study was funded by core funding of the UK Medical Research Council (MRC) to the MRC Human Genetics Unit. We are indebted to the ICGC for the timely public release of tumour whole genome sequencing data.

## AUTHOR CONTRIBUTIONS

The study was conceived and the manuscript written by V.B.K. and C.A.S. The data was analyzed by V.B.K.

## SUPPLEMENTAL FIGURE LEGENDS

**Supplemental Figure S1:** DSB rates in the NHEK cell line in the vicinity of NHEK LAPs. DSB rates were measured in the NHEK cell line within 500 Kb of NHEK LAPs, centred on the CTCF binding motif of each LAP. DSB rates are shown for LAPs that overlap ENCODE chromHMM annotated NHEK (A) promoters (762 LAPs), (B) enhancers (1,485 LAPs), or (C) neither promoters or enhancers (2,030 LAPs).

**Supplemental Figure S2:** Germline mutation rates contrast with somatic mutation rates at LAPs. (A) DNA breakpoints in the ICGC pan-cancer analysis increase at LAPs but not at recombination HSs. (B) *De novo* germline variants from the DECODE trios dataset show no increase at LAPs or HSs. Light blue: genomic regions centred around recombination HSs; dark blue: centred around LAPs.

**Supplemental Figure S3:** Variation in chromatin features around LAPs and HSs. Average overlap per Kb with HepG2 cell line ChIP-seq peaks around LAPs and HSs for (A) H3K9me3, (B) H3K27ac, (C) H3K27me3, (D) H3K36me3, (E) H3K4me3. (F) Average replication timing. (G) Average overlap per Kb with HepG2 DNAse HS sites; (H) Average GC content; (I) Average genic sequence per Kb; (J) Average overlap per Kb with MCF-7 cell line CTCF ChIP-seq peaks. Light blue: centred around recombination HSs; dark blue: centred around LAPs.

**Supplemental Figure S4:** Replication timing varies among functional sub-regions within loop anchor points. LAPs of the HMEC cell line stratified by chromHMM HMEC predicted chromatin states (promoters, enhancers, insulators), relative to their replication timing in MCF-7 cells.

**Supplemental Figure S5:** Mutational landscapes vary among tumour types. (A) SV Breakpoint rates within the vicinity of LAPs stratified by cancer type. (B) SNV rates at LAPs stratified by cancer type.

**Supplemental Figure S6:** Mutational spectra in BRCA-deficient tumours. (A) Mutational spectra observed for BRCA1/2 wild-type (++) (top two rows) and BRCA1/2 deficient (−−) (bottom two rows) samples at LAPs and control sites located 1 Mb away from the LAPs. (B) The decrease in RSS (residual sum of squares) and the increase in explained variance with the addition of more signatures to the mutational model. (C) Mutational signature heatmap, showing the relative enrichment and depletion of mutations in their trinucleotide context (y-axis), stratified by mutational signature (x-axis). (D) The association of mutational signatures with LAPs and control sites (LAPs+1MB) for BRCA1/2 wild-type (++) and BRCA1/2 deficient (−−) samples.

**Supplemental Figure S7:** LAP discovery is a function of Hi-C sequencing depth.

The number of private, i.e. cell line-specific, CTCF binding site containing LAPs plotted against the total number of LAPs called in each cell line assayed by Rao et al. (2014).

